# Nutritional composition and metabolic analysis of the newly hatched *Anguilla japonica* from embryo to preleptocephali obtained from artificial reproduction

**DOI:** 10.1101/2023.05.13.539677

**Authors:** Kang Li, Yuangu Li, Tiezhu Li, Rongfeng Cui, Liping Liu

## Abstract

The starter diet for Japanese eel (*Anguilla japonica*) has always been a difficult problem for the realization of total artificial reproduction. Therefore, this research analyzed the nutritional composition of artificially fertilized eggs, and transcriptome of samples from early hatchlings of fry to better understand nutrients requirements. The composition of crude lipid and crude protein in fertilized eggs was 7.24%±0.32% and 10.56%±0.41%, respectively. Seven kinds of essential amino acids (EAA) were detected but took a comparable lower content (3.19%) than other marine fish eggs. We randomly assembled 265.74 million clean reads and identified 1751 differentially expressed genes (DEGs) (*P* < 0.01) from pre-leptocephalus larvae. A total of 23 KEGG pathways related to the digestive and metabolic system were detected. Genes related to the secretion pathway of saliva, pancreatic juice and other digestive juices were significantly changed. The genes of carbohydrate metabolism, glycerolipid metabolism and glycerophospholipid metabolism were up-regulated significantly with the growth of the larvae (day 0 to 12). This study will facilitate future studies on the nutrition of eel larvae and other biological traits to reproductive research.

## 1. Introduction

Japanese eel (*Anguilla japonica*) is an important cultured species in East Asian countries with a mystery life cycle involving spawning in the ocean before returning to freshwater (Kuan-Mei, et al., 2018; Tanaka, 2015; Tsukamoto, 1992). Its populations have suffered a severe reduction due to intensive exploitation, habitat loss, and environmental deterioration (Jacoby and Gollock, 2014; Matsushige, et al., 2019). Farming of eels, which depends on wild-caught glass eel, is also experiencing a critical decrease in the annual harvest (Dekker and Casselman, 2014; Koh, et al., 2017; Miller, et al., 2009). The complete life cycle of *A. japonica* was firstly closed in the lab in 2010 (Masuda, et al., 2012; Tanaka, et al., 2001). However, the commercialization of lab-produced larvae is still hindered by the low survival rate of pre-leptocephalus larvae (Okamura, et al., 2014).

Low survival rate in lab is likely due to nutritional deficiencies in their artificial diets, despite previous efforts to provide diets including shark egg yolk, Antarctic krill, and rotifer paste, additional nutritional improvements are still needed (Liu, et al., 2017; Okamura, et al., 2019; Okamura, et al., 2013; Tanaka, et al., 2003). Pousão-Ferreira, et al. (1999) have used gilthead seabream (*Sparus aurata*) eggs to feed *S. aurata* larvae successfully and determined its nutritional requirements during the larval rearing based on the nutritional composition of *S. aurata* eggs. Moreover, Mourente and Rosa (1996) confirmed the fatty acids levels of feed for unfed larvae of the Senegal sole (*Solea senegalensis*) according to their egg’s composition. The changes of major yolk nutrients from fertilized eggs to yolk-sac larvae of Japanese eel described by Ohkubo, et al. (2008) emphasizes the importance of understanding egg nutritional composition for developing the starter diets.

Digestive system function of fish larvae relies on the onset of genetically pre-programmed and extrinsically influenced factors (Politis, et al., 2018). The expression of several selected pancreatic enzyme genes indicated that *A. japonica* larvae acquire full function by the onset of exogenous feeding at 8 dph (Kurokawa, et al., 2002). However, other digestive enzyme genes and nutrient transporters of *A. japonica* are not well studied, and few studies have addressed the food digestion and nutrient absorption during the preleptocephali stage. In recent years, a new generation of sequencing technology has been widely applied to the genome and transcriptome analysis of aquatic organisms (Kleppe, et al., 2014). Changes in the quantity and quality of transcriptome data can reflect the expression status of related genes in specific circumstances, which can effectively improve the quality of basic research. RNA-Seq has been widely used as an effective tool for transcriptome analysis to discover, profile, and quantify the RNA transcripts (Wang, et al., 2010). It provides fundamental insights into biological processes and applications such as gene expression levels in developmental stages, combining advantages of high throughput, low background noise and high sensitivity (Churcher, et al., 2015; Hsu, et al., 2015; Ozsolak and Milos, 2010).

With the aim to explore the dietary requirements for the preleptocephali of *A. japonica* during the critical early life history stage, the nutritional composition of eggs was determined and the RNA of both embryos and preleptocephali was sequenced. It is hope to reveal the molecular events that affect the nutrient digestion and absorption, and to provide a basis for the future research on nutrient requirements of artificially eel hatchlings.

## 2. Materials and Methods

### 2.1. Artificial reproduction and sampling

The wild-caught broodstock Japanese eels were cultured in the experimental facility of the Marine and Fisheries Research Institute of Ningbo (China), where they were kept in 8 m × 4 m × 1.7 m pond and the natural seawater was maintained at ∼ 20 °C and adjusted to ∼ 31 psu salinity using Red Sea® Salts. Fish were not fed during the experiment and around 75% of the pond area was shaded by the black net. At the onset of experiments, all experimental broodstock eels were anaesthetized (MS-222) and then tagged with a passive integrated transponder.

The Japanese eels were artificially induced to mature using the method detailed in Jiang, et al. (2012). Five female broodstocks were euthanized after stripping to collect eggs, which were pooled at equal ratios before being stored at -80 °C until nutritional analyses. Natural spawning happened around 12 h after the remaining females and males were put together, then fertilized eggs were collected and reared at 24 °C in seawater (31 psu) in darkness. Hence, 50 fertilized eggs were collected as 0 d sample, and 30 pre-leptocephalus samples were taken every three days after hatching until the 12^th^ day. All samples were preserved by RNA*later* (Tiangen Biotech, Beijing, China) and then stored at -80 °C until RNA extraction.

### 2.2. Nutritional analysis of eggs

The proximate composition analysis of eggs was performed using official methods of analysis of AOAC (1996). Moisture content was determined by drying eggs’ sample in an oven at 110 °C to a constant weight. Ash content of eggs was determined by heating the sample in a muffle furnace for about 10-12 h at 550 °C and weighting it after cooling. Crude protein content was measured by the Kjeldahl method (N content × 6.25) using a Kjeltech system (Kjeltec 2300, Foss, Sweden). The extraction of lipids from samples was carried out with a mixture of chloroform and methanol (2:1, v/v) containing 0.01% BHT and determination of crude lipids was performed according to Folch, et al. (1957).

Fatty acid methyl esters (FAMEs) were esterified with 14% BF_3_ in methanol and the FAMEs extracted with hexane (Morrison and Smith, 1964). FAMEs were separated and detected by an Agilent 7890 gas chromatograph (Agilent Technologies, CA, USA) equipped with a flame ionization detector instrument. The 37-FAME Mix (Supelco, Bellefonte, PA, USA) was used to identify the FAMEs, and the fatty acid C19:0 (nonadecanoic acid, Sigma) was used as an internal standard for fatty acid quantification. All measurements were performed in triplicates, the fatty acids content was expressed as the percentage of each FA to the total FAs. The amino acid composition of the freeze-dried eggs digested with hydrochloric acid was determined with a HITACHI L-8900 amino acid automatic analyser (Hitachi Limited, Tokyo, Japan), and the peak areas were recorded (Gilani and Peace, 2005). Standard curves were plotted with amino acid peak area as the ordinate and amino acid concentration as the abscissa.

### 2.3. Calculation of the theoretical demand of nutrients in the starter diets of larvae eels

The theoretical demand of each nutrient was calculated according to previous research (Li, et al., 2016; Pousão-Ferreira, et al., 1999). The crude lipid and crude protein contents of the fresh weight base were converted into dry weight base, which was the theoretical requirement for fat and protein of larvae eels. The requirement of each amino acid can be converted according to the content of crude protein on the basis of dry weight. Since the amino acids in this study were tested on the basis of dry weight, the detected value is the theoretical requirement of amino acids in the feed of larvae eels. Similarly, according to the crude fat content based on dry weight, the theoretical requirement of fatty acids in the feed can be calculated.

### 2.4. RNA extraction, library construction and sequencing

The total RNA of the whole fish was extracted using TRIzol® reagent (Invitrogen, California, USA) according to the manufacturer’s instructions. Purified RNA was quantified by Agilent 2100 Bioanalyzer (Agilent Technologies, CA, USA). From each pooled sample, 5μg mRNA was isolated from total RNA using oligo (dT) magnetic beads (Invitrogen). Then five sequencing libraries, one for each time point, were constructed using Truseq™ RNA sample prep Kit (Illumina, San Diego, USA) according to the instruction. The mRNA was interrupted into∼200 bp short fragments using the fragmentation buffer. It was then transcribed into the first-strand cDNA using reverse transcriptase and random hexamer primers, followed by second-strand cDNA synthesis. The double-stranded cDNA was subjected to end repair, phosphorylation, a-tailed and indexed adapters were ligated. Suitable fragments were selected and enriched by PCR to create the final cDNA library. The paired-end cDNA library was sequenced on an Illumina HiSeq™ 4000 platform (Majorbio Biotech Co., Ltd., Shanghai, China).

### 2.5. De novo *assembly and functional annotation*

The raw RNA-seq data were processed to discard the dirty reads that include reads with adaptors, reads with more than 10% Q<20 bases. The low complexity reads were removed by Seqprep and Sickle program. Clean and high-quality reads from the five samples were then assembled using the Trinity program, meanwhile, the counts of transcripts and the N50 were calculated. The assembled unigenes of five samples were used for BlastX search and annotation against the NR, Swissprot, COG (Clusters of Orthologous Groups of proteins), KEGG (Kyoto Encyclopedia of Genes and Genomes) and GO (Gene Ontology) databases with a cut-off E-value < 10^−5^. Based on Nr-matched unigenes, the annotation of GO was obtained by Web Gene Ontology Annotation Plot (WEGO; http://wego.genomics.org.cn/) program. The transcripts were also blasted against the Pfam database to identify specific protein domains and acquire GO annotations.

### 2.6. Comparative expression analysis

All clean sequence reads from each of the five libraries (0, 3, 6, 9, 12 days) were mapped to reference sequences (unigenes from the assembled transcriptome data) using Bowtie2 software with default setting (Langmead, 2012). Subsequently, RSEM (http://deweylab.biostat.wisc.edu/rsem/) was used to calculate the FPKM (Fragments per kilobase of transcripts per million fragments mapped values) of the assembled transcripts (Li and Dewey, 2011). Identifying differentially expressed genes (DEGs) among five groups was performed using the R package WGCNA (Robinson, et al., 2010). Because we have a single sample from each time point, pair-wise comparisons between time points are not feasible. We decided to examine time-dependent transcriptional changes using a linear model where time points are the independent variable and expression levels are the dependent variable. The false discovery rate (FDR) method was introduced to determine the threshold *P*-value in multiple tests. If FDR was smaller than 0.05 and FPKM values showing at least twofold difference two groups, this unigene was considered as significant time-dependent DEGs. DEGs among the samples were further annotated by GO and KEGG pathway analysis.

### 2.7. Statistical analysis

In the experiment, the data of each group were presented as mean ± standard deviation (SD) and analyzed by Excel 2010 statistical software and SPSS version 17 (Michigan Avenue, Chicago, IL, USA). The original transcriptomic data were analyzed by a linear model implemented in the WGCNA package. With day-age as the independent variable and gene expression as the dependent variable, genes significantly changed with day-age were identified. We applied a WGCNA module significance filter of *P*< 0.01 and DEG FDR < 0.05 for the final DEG gene set.

## 3. Results

### 3.1 Nutrition composition of eggs

The published fatty acid composition of different marine fish eggs was summarized in Table 1.

**Table 1.**
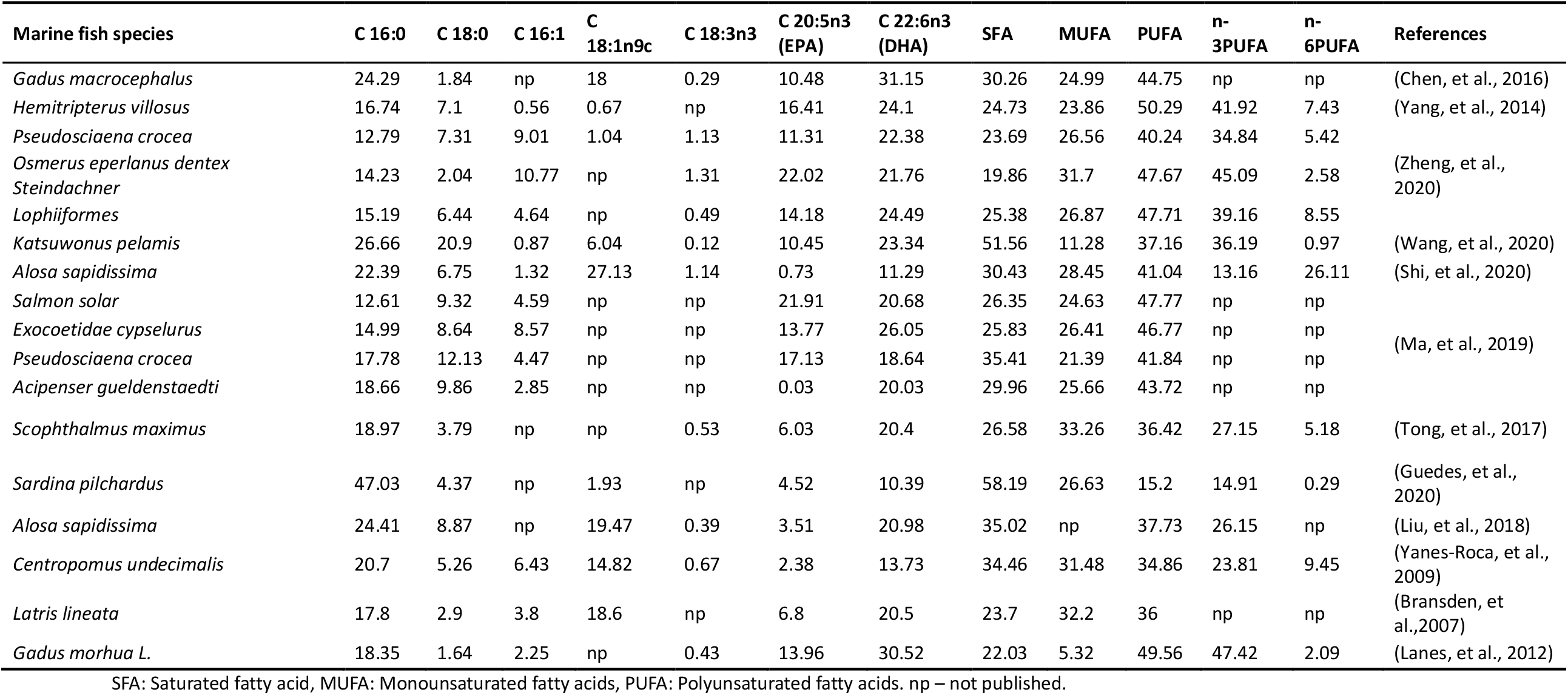
Analysis of fatty acid composition of different marine fish eggs (% total fatty acid).

Proximate composition of unfertilized eggs is shown in Fig. 1. The composition of crude lipid and crude protein was 7.24%±0.32% and 10.56%±0.41%, respectively. A total of nine saturated fatty acids (SFA) were detected in the eggs, together with 8 monounsaturated fatty acids (MUFA), and 10 polyunsaturated fatty acids (PUFA), which took 41.14%±1.61%, 18.19%±0.69%, and 40.69%±1.11% of total fatty acids. The n-3 PUFA takes up about 34.03%±0.50% of total fatty acids, while the n-6 PUFA took around 5.04%±0.48% in total. The EPA (C20:5n3) and DHA (C22:6n3) accounted for 4.27%±0.24% and 25.98%±0.49% of total fatty acids, respectively, in which EPA was lower than most marine fish, and DHA was higher than most marine fish (Fig. 1 and Table 1). Moreover, the total EPA and DHA content is 30.25% is medium level, comparing to other marine fish. The SFA of eel eggs accounted for 41.14%±1.61% of the total fatty acids, which was higher than most marine fishes listed in Table 1, among which the content of C16:0 was the highest. Simultaneously, compared to the contents of fatty acids in other marine fish eggs, it was found that eel eggs were rich in fatty acids, which provided a good foundation for the development of eel larvae (Table 1 and Fig. 1). The theoretical fat demand of starter diet was 40.86% in the current study, that the demand of C16:0 and DHA was the highest (Table 2).

**Table 2.** Theoretical requirements of nutrients of *A. japonica* larvae (%, dry weight).

| Items | Theoretical requirement | Items | Theoretical requirement |
| --- | --- | --- | --- |
| Protein | 59.59 | Fat | 40.86 |
| Glutamate | 0.98 | C16: 0 | 12.89 |
| Cysteine | 0.16 | C20: 4n6 | 1.24 |
| Leucine | 0.69 | C20: 5n3 | 1.87 |
| Lysine | 0.63 | C22: 6n3 | 11.39 |
| Alanine | 0.82 | n-3 PUFA | 14.91 |
| Valine | 0.42 | n-6 PUFA | 2.21 |

**Fig. 1.**
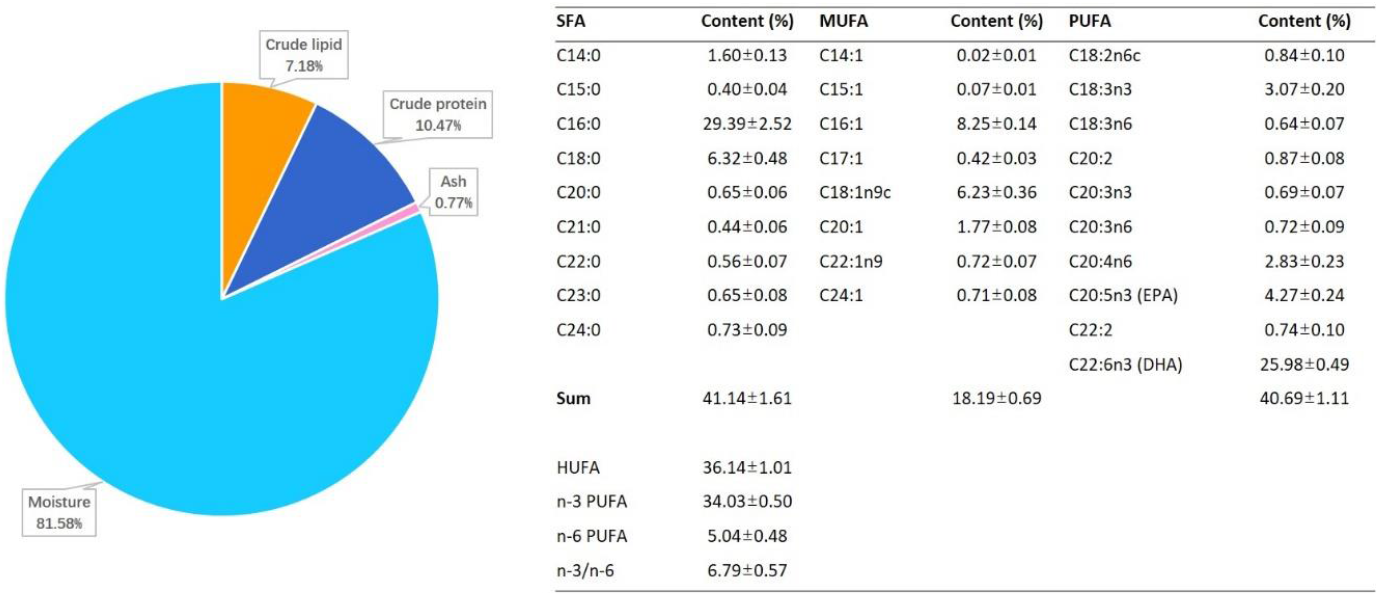
The proximate composition (% wet weight basis) and fatty acid composition (% total FA) in unfertilized eggs of *Anguilla japonica*.

A total of 17 amino acids were detected, including 7 kinds of essential amino acids (EAA) and 10 nonessential amino acids (NEAA, Fig. 2). Leucine (0.69%) took a higher percentage compare to other EAAs, and the glutamic acid was the highest (0.98%) among NEAAs. The ratio of EAA/NEAA was 65.37%, and the EAA made up about 39.53% of the total amino acids (Fig. 2). Compared with other amino acids, the contents of glutamic acid (0.98%), alanine (0.82%), leucine (0.69%), and lysine (0.63%) in artificial breeding eel eggs are higher, and glutamate is the highest, which is similar to most marine fish. Still, the content of each amino acid is far lower than that of other marine fish eggs (Table 3). The theoretical demand for protein in the diet of the larvae was 59.59%, the theoretical demand for glutamate was the highest, and the theoretical demand for cysteine was the lowest (Table 2).

**Table 3.**
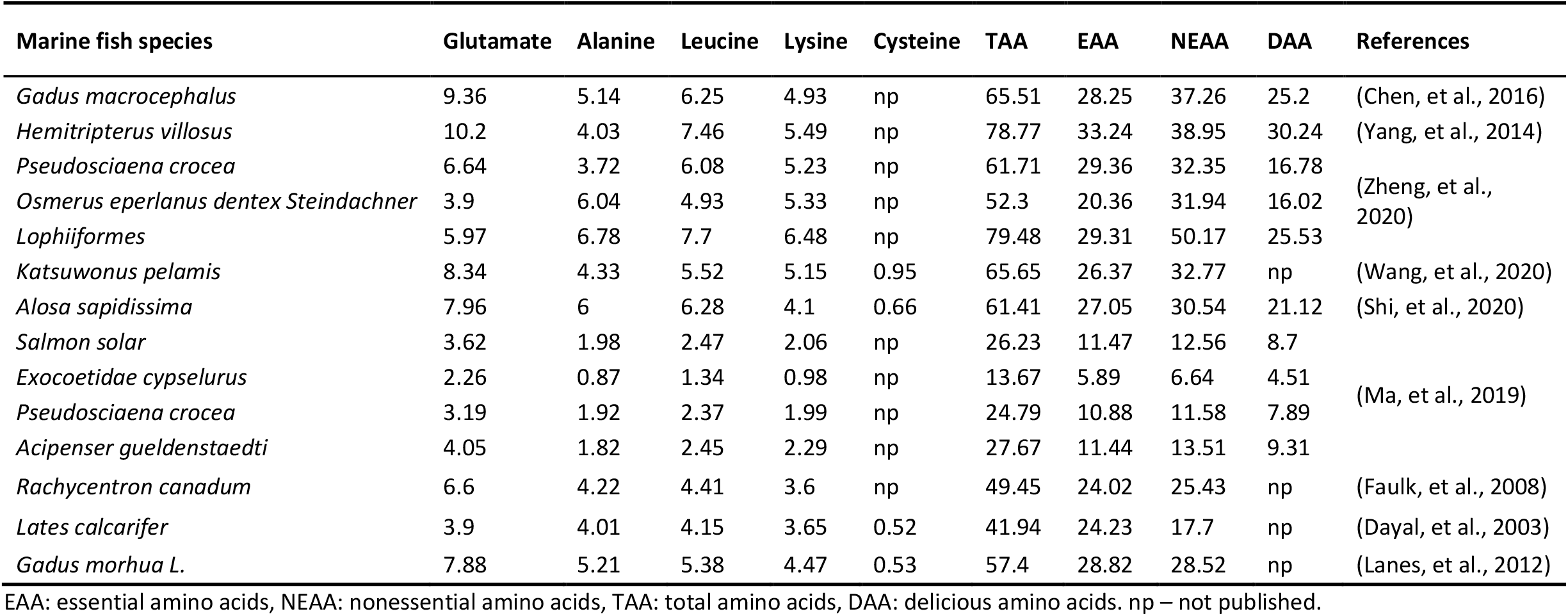
Analysis of amino acid composition of different marine fish eggs (% dry weight basis).

**Fig. 2.**
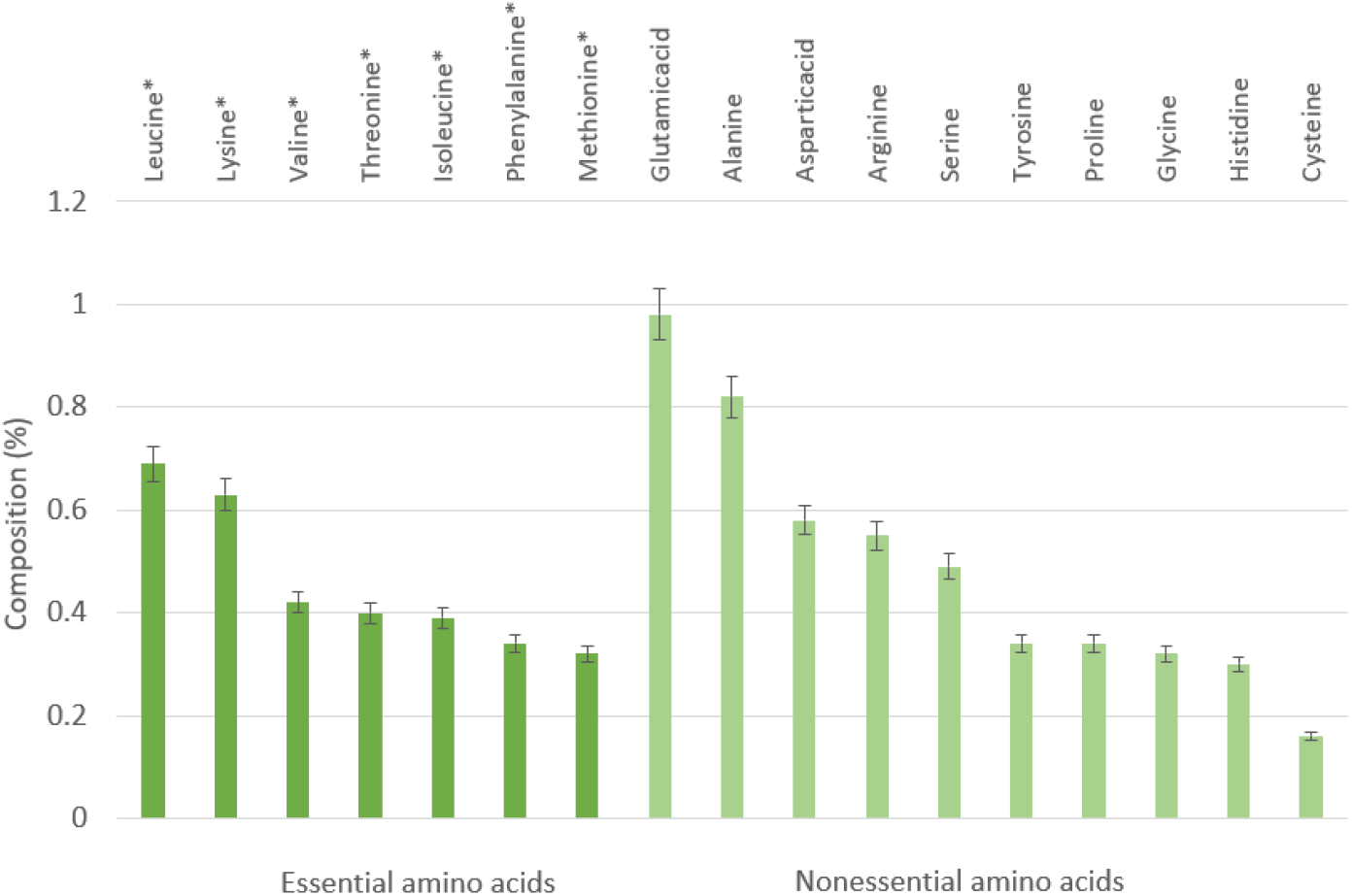
Composition of essential amino acids and nonessential amino acids in unfertilized eggs of *A. japonica*.

### 3.2 Processing of sequencing data and de novo assembly

The RNA sequencing generated a total of 271,423,952 raw reads, and 265,743,784 (97.9%) clean reads were obtained after removing SeqPrep adapter and low-quality reads (Table 4). Thereafter, we got a total of 102941 unigenes with a Q20 percentage over 98%. The contig N50 was 1398 bps. The average length of it was 856.6 bps (Table 5). For the functional annotation, the unigenes were aligned with sequences from major databases including Pfam, Swiss-Prot, KEGG, GO and Nr. The statistics of overall functional annotations were shown in Table 6.

**Table 4.** Read data of *Anguilla japonica* before and after processing.

| Day-age | Number of raw read pairs | Number of cleaned read pairs |
| --- | --- | --- |
| 0 | 49,215,850 | 48,233,440 (98.0%) |
| 3 | 54,441,304 | 53,379,346 (98.0%) |
| 6 | 58,638,172 | 57,402,690 (97.9%) |
| 9 | 51,574,270 | 50,400,978 (97.7%) |
| 12 | 57,554,356 | 56,327,330 (97.9%) |

**Table 5.** Statistics of assembly of *A. japonica*.

| Type | Unigene |
| --- | --- |
| Total sequence number | 102,941 |
| Total sequence base | 88,179,496 |
| Percent GC | 54.92% |
| Largest length | 26610 bp |
| Smallest length | 201 bp |
| Average length | 856.6 bp |
| N50 | 1398 bp |

**Table 6.** Statistics of unigene functional annotation.

| Database | Annotated unigenes | Percentage |
| --- | --- | --- |
| Pfam | 30993 | 30.11% |
| KEGG | 33072 | 32.13% |
| GO | 25245 | 24.52% |
| Swiss-prot | 45067 | 43.78% |
| Nr | 56574 | 54.96% |

### 3.3 Identification and analysis of the differently expressed genes (DEGs)

A summary of unigenes classified to each term at GO level 2 is shown in Fig. 3. A total of 1664 terms in the biological process category were produced, the most dominant subcategories were cellular process (17087; 21.14%), single-organism process (12101; 14.97%), metabolic process (11794; 14.59%), biological regulation (7441; 9.21%), and regulation of biological process (6915; 8.56%). In cellular component functions, we got 774 terms. And there were 15712 (19.89%) of unigenes that were assigned to the cell, followed by cell part 15469 (19.58%), membrane 11852 (15.00%), organelle 9858 (12.48%), and macromolecular complex 5782 (7.32%). In the molecular function category, 422 terms were produced, and the five most dominant subcategories were binding (20754; 43.91%), catalytic activity (15442; 32.67%), transporter activity (2474; 5.23%), signal transducer activity (2054; 4.35%) and structural molecule activity (1971; 4.17%).

**Fig. 3.**
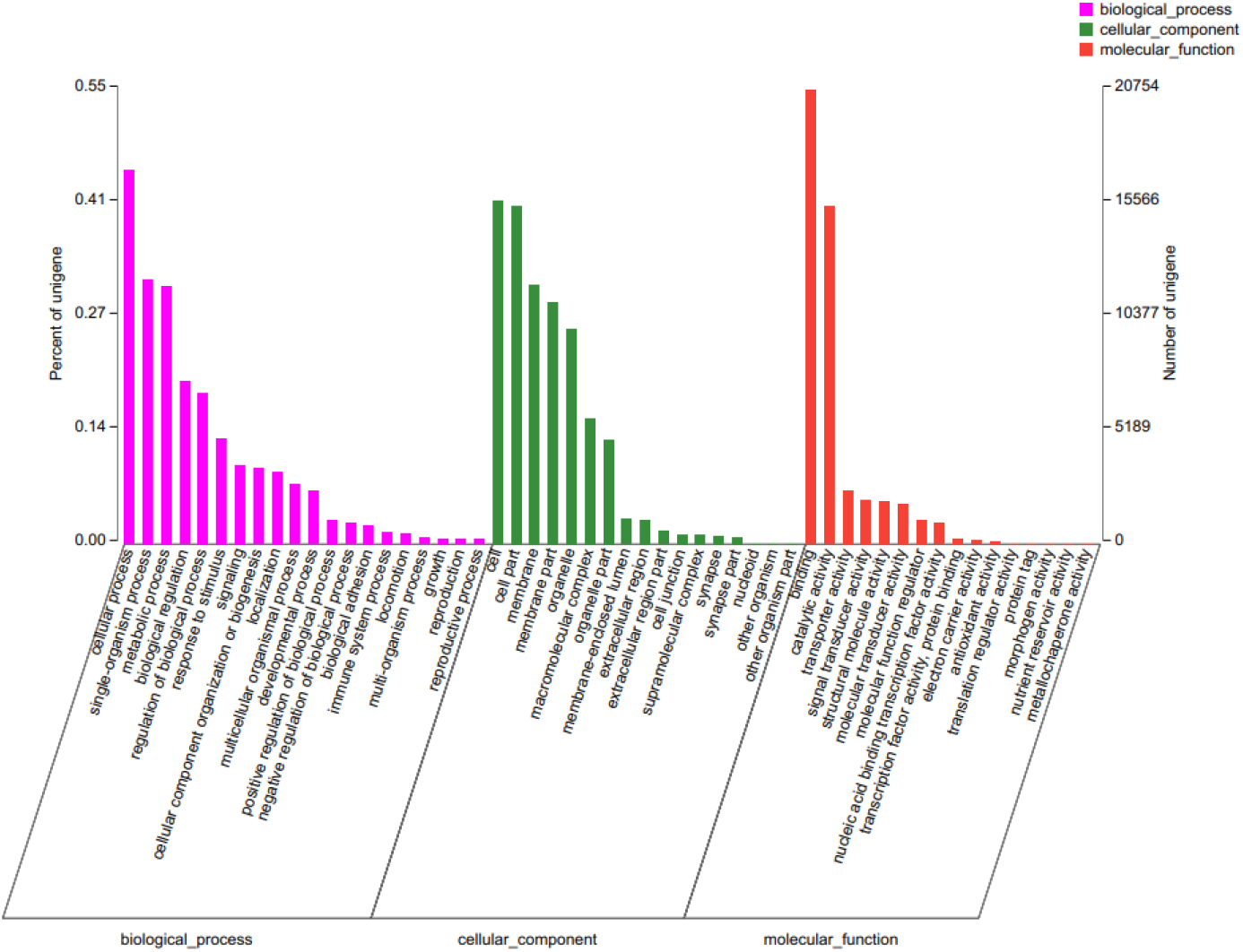
Functional annotation of assembled contigs associated with GO terms.

By running the WGCMA to perform a linear regression of time-dependent gene expression on the original transcriptome data, genes significantly changed with day-age were identified. A total of 1,751 genes were selected that changed significantly with age, contained in WGCNA module with *P* < 0.01 and gene DEG FDR < 0.05. By comparing and analyzing the screened genes with KEGG database, 23 pathways related to the digestive and metabolic system were detected. With the change of age, 120 genes showed a significant increase in gene expression, and 18 genes showed a significant decrease in gene expression. Among them, genes related to the secretion pathway of saliva, pancreatic juice and other digestive juices were significantly changed. The genes of carbohydrate metabolism, glycerolipid metabolism, glycerophospholipid metabolism, and other metabolic pathways were up-regulated significantly with age (Table 7).

**Table 7.** Digestive and metabolism-related pathways in KEGG term.

| Pathway id | Pathway | Up No. | Down No. | Total No. |
| --- | --- | --- | --- | --- |
| Map04972 | Pancreatic secretion | 12 | 1 | 13 |
| Map04970 | Salivary secretion | 13 | 1 | 14 |
| Map04971 | Gastric acid secretion | 10 | 1 | 11 |
| Map04976 | Bile secretion | 4 | 1 | 5 |
| Map04973 | Carbohydrate digestion and absorption | 7 | 0 | 7 |
| Map04974 | Protein digestion and absorption | 4 | 0 | 4 |
| Map04975 | Fat digestion and absorption | 2 | 0 | 2 |
| Map04978 | Mineral absorption | 1 | 0 | 1 |
| Map04977 | Vitamin digestion and absorption | 0 | 0 | 0 |
| Map00310 | Lysine degradation | 6 | 3 | 9 |
| Map00250 | Alanine, aspartate and glutamate metabolism | 1 | 2 | 3 |
| Map00380 | Tryptophan metabolism | 2 | 0 | 2 |
| Map00350 | Tyrosine metabolism | 2 | 0 | 2 |
| Map00270 | Cysteine and methionine metabolism | 0 | 1 | 1 |
| Map00600 | Sphingolipid metabolism | 5 | 1 | 6 |
| Map01040 | Biosynthesis of unsaturated fatty acids | 2 | 0 | 2 |
| Map00561 | Glycerolipid metabolism | 11 | 1 | 12 |
| Map00564 | Glycerophospholipid metabolism | 13 | 1 | 14 |
| Map00562 | Inositol phosphate metabolism | 9 | 3 | 12 |
| Map00051 | Fructose and mannose metabolism | 5 | 1 | 6 |
| Map00052 | Galactose metabolism | 4 | 1 | 5 |
| Map00500 | Starch and sucrose metabolism | 5 | 0 | 5 |
| Map00010 | Glycolysis / Gluconeogenesis | 3 | 0 | 3 |

Statistical overrepresentation test was performed for genes whose gene expression significantly increased with age after screening. A total of 20 significantly enriched Go-slim molecular function pathways were detected. Through analysis, it is found that among the genes enriched in the transmembrane transporter activity pathway, the TRPC1 gene and CFTR gene are related to the secretion and transport of digestive enzymes such as saliva, pancreatic juice, and bile (Table 8).

**Table 8.** Enrichment of genes significantly different with age in GO term

| Pathway id | Pathway | Number of matched GO | +/- | FDR |
| --- | --- | --- | --- | --- |
| GO:0005262 | calcium channel activity | 7 | + | 0.016900 |
| GO:0015085 | calcium ion transmembrane transporter activity | 7 | + | 0.015700 |
| GO:0046873 | metal ion transmembrane transporter activity | 19 | + | 0.000202 |
| GO:0022890 | inorganic cation transmembrane transporter activity | 19 | + | 0.000737 |
| GO:0008324 | cation transmembrane transporter activity | 22 | + | 0.000092 |
| GO:0015075 | ion transmembrane transporter activity | 26 | + | 0.000103 |
| GO:0022857 | transmembrane transporter activity | 31 | + | 0.000032 |
| GO:0005215 | transporter activity | 33 | + | 0.000031 |
| GO:0015318 | inorganic molecular entity transmembrane transporter activity | 25 | + | 0.000128 |
| GO:0005261 | cation channel activity | 13 | + | 0.007660 |
| GO:0005216 | ion channel activity | 16 | + | 0.001290 |
| GO:0015267 | channel activity | 17 | + | 0.000836 |
| GO:0022803 | passive transmembrane transporter activity | 17 | + | 0.000940 |
| GO:0015081 | sodium ion transmembrane transporter activity | 8 | + | 0.008010 |
| GO:0015077 | monovalent inorganic cation transmembrane transporter activity | 14 | + | 0.001780 |
| GO:0015079 | potassium ion transmembrane transporter activity | 10 | + | 0.019700 |
| GO:0008509 | anion transmembrane transporter activity | 11 | + | 0.025100 |
| GO:0004930 | G protein-coupled receptor activity | 11 | + | 0.040200 |
| GO:0004888 | transmembrane signaling receptor activity | 16 | + | 0.028700 |
| GO:0038023 | signaling receptor activity | 18 | + | 0.027100 |

## 4. Discussion

Studies have demonstrated that the knowledge of nutrient composition of fish eggs will not only provide key insights into requirements of nutrient, but also support formulating the starter diet (Pousão-Ferreira, et al., 1999; Yang, et al., 2014). Meanwhile, the transcriptomic analysis could reveal the changes of food digestion and nutrient absorption during the preleptocephali stage to eliminating the hurdle to complete full artificial reproduction.

As an essential component of cells and tissues, protein is an essential substance for animal growth and life maintenance (Zhou, et al., 2014), furthermore, the obtained data of egg’s protein content emphasize the nutritional requirements of the first larval feeding (Pousão-Ferreira, et al., 1999). The theoretical protein demand of starter diet of Japanese eel was 59.59% that calculated depending on the crude protein composition (10.56%) of eel eggs in present study, which is consistent with the theoretical value of glass eel studied by Xiong, et al. (1996). However, the difference from this study is that EAA content (3.19%) was far lower than the eggs of other marine fish, hence it might be a cause that led to malnutrition and mass death before feeding. Known as the first restrictive amino acid, lysine could improve the utilization of other EAAs and prevent nitrogen loss, as well as promote fish development (Zhou, et al., 2008). Therefore, a higher lysine content in fish egg is beneficial to both fish growth and survival. Although the content of lysine (0.69%) was one of the highest amino acid in the eel eggs of this study, a comparable lower rate to other marine fish eggs still implies that a limited lysine content could be another cause for the mass death in the artificial reproduction. Fatty acids are one of the main energy sources for fish (Bennett, et al., 2007; Cejas, et al., 2004). The analysis shows that the SFA of *A. japonica* eggs accounts for 41.14% of the total fatty acids, among which the content of C16:0 was the richest that is similar to *Alosa sapidissima* (Yanes-Roca, et al., 2009), *Sardina pilchardus* (Liu, et al., 2018), and *Centropomus undecimalis* (Guedes, et al., 2020). PUFA has been proved to have a significant effect in promoting fish growth and development, as well as the fish immunity and survival rate (Youqing, et al., 2010). A 40.69% of the total fatty acids’ PUFA was detected in this study, and a similar high proportion was found in other marine fish eggs as well (Lin, et al., 2020; Liu, et al., 2018; Zheng, et al., 2020).

In this study, the transcription levels of various digestive enzymes and carbohydrate metabolism in the larvae were comparably high with the changes of the larvae’s age. The results of this study indicated that the metabolic capability of low molecular carbohydrate kept increasing with the day-age growing, especially in the pathways of galactose, fructose, and sucrose metabolism. The hyaluronic acid, which is a disaccharide substance, is found to be the main body composition of leptocephali, indicates that carbohydrate is an critical substance for the growth of leptocephali (Pfeiler, 1999). The sea snow, a most likely starter diet of Japanese eel, happens to be a collection of different carbohydrates also supports the demands of low molecular carbohydrate during the early life (Pfeiler, 1999). Okamura, et al. (2020) found that dietary supplementation with chitin hydrolysates including mono-, di- and trimers of N-acetylglucosamine had positive effects on the growth and survival of eel larvae. It demonstrates that feeding low-molecular bait such as glucose sugar and maltose to larvae eel is beneficial to improve the survival time of larvae (Cowen, et al., 1996; Skoog, et al., 2008), which is consistent with our experimental results. During the early stage of post-hatching, the larvae consume endogenous yolk protein to supply the growth needs (Nobuyuki et al., 2008). It implies that the consumption of yolk nutrient continued after the larvae hatch, and the body carbohydrates kept strengthen in order to meet the energy demand for its growth. Our previous study indicated that the teeth began to form on the 6^th^ day of membrane emergence, and the oil globule completely disappeared from the day 8, which marked the transition from endogenous nutrition to exogenous nutrition of larvae eel (Liu et al., 2017).

Amino acids play essential role in fish metabolism. On the one hand, amino acids can be used as signal molecules to participate in various physiological regulation of the body. On the other hand, in the case of hunger or malnutrition, amino acids can also be oxidized to provide energy through gluconeogenesis (Ca, et al., 2004; R?Nnestad, et al., 1999). Transcriptome analysis showed that the metabolic transcription level of lysine increased steadily with the change of age. The lysine content in fish eggs was relatively high that could meet the needs of early growth of larva, and lysine is also one of the important ketogenic amino acids. When there is a lack of available energy in the organism, lysine can participate in the generation of ketone body and glucose metabolism, and it is an vital energy material supplement in the organism (Huang, et al., 2021).

Transcriptome analysis showed that with the growth of age, the level of glycolytic metabolism in larvae increased significantly (Table 7), and the transcription level of hexokinase (HK), which is involved in the catalytic glycolytic metabolism pathway, also increased notably. This indicates that in order to meet the growth nutritional requirements of the larvae, glucose and other sugars in the body should be converted into energy to meet the growth needs. At the same time, it was found that the transcription level of gluconeogenesis in larvae increased with the change of age. The metabolism and transcription levels of glyceride and glycerophospholipid were enhanced. This suggests that during the growth process, larvae may have insufficient carbohydrate energy reserves and demands to convert other non-carbohydrate substances into glucose or glycogen to meet the energy supply needed for growth.

A previous study presented that the growth rate of the farmed leptocephali (which feeds on shark eggs) is lower than that of the wild leptocephali (Ishikawa, et al., 2001). As one of the commonly used artificial starter diet components, the *Acanthias* shark egg-based diet are composed with protein (26.3%), lipids (17.5%), carbohydrates (0.1%) and moisture (54.4%) (Okamura, et al., 2014). Our results indicated that the carbohydrate conversion in larvae increased with the increase of the age of larvae. We hypothesized that the slow growth rate of the cultured preleptocephali larvae might be related to the insufficient addition of carbohydrate in the bait. Okamura, et al. (2020) presented that dietary supplementation with high sugar content such as N-acetylglucosamine, glucose and maltose could significantly improve the growth rate of larvae. Moreover, this study indicates that the egg contains enough fatty acids to meet the nutritional demands of the preleptocephali stage. According to the results of this study, it suggests that the proportion of low molecular carbohydrates and the content of EAA such as lysine can be increased appropriately in the feeding of artificially bred eel larvae, and the specific mechanism of digestion and metabolism needs further experimental verification. Among the functional pathways of significantly enriched GO-Slim molecules, most of the pathways are related to ion transport, which may be related to early nutrient metabolism and transport, and might be related to water salinity, which requires further analysis.

## 5. Conclusions

In present study, the nutritional composition and transcriptome of the artificially fertilized eggs and pre-leptocephalus larvae were investigated. Seven kinds of essential amino acids (EAA) were detected, and the results presents a lower content in *A. japonica* than other marine fish eggs. A total of 265.74 million clean reads were randomly assembled and 1751 differentially expressed genes were identified. The genes of carbohydrate metabolism, glycerolipid metabolism and glycerophospholipid metabolism were up-regulated significantly with the growth of the larvae. This result provides a theoretical basis for future studies on the nutrient requirements and digestive function of the newly hatched *A. japonica*, promotes the overall artificial reproduction of eel.

## Data availability

The data generated and/or analyzed in the current study are provided in the paper.

## Ethical statements

All experiments involving animals were approved by the Animal Care and Use Ethics Committee of the Shanghai Ocean University.

## Author contributions

**Kang Li and Yuangu Li**: conceptualization, methodology, investigation and writing; **Tiezhu Li**: investigation, methodology, and writing; **Rongfeng Cui:** methodology, and writing-review; **Liping Liu**: writing-review & editing, supervision and funding acquisition. All authors read and approved the final manuscript.

## References

AOAC, 1996. Official methods of analysis of the association of official analytical Chemists. Arlington (U.S.A.): Association of Official Analytical Chemists.

Bennett, P.M., Weber, L.P., Janz, D.M., 2007. Comparison of chloroform–methanol-extracted and solvent-free triglyceride determinations in four fish species. J Aquat Anim Health. 19, 179–185.

Ca, A., Lecc, A., Mtd, A., Hjf, B., 2004. Amino acid pools of rotifers and Artemia under different conditions: nutritional implications for fish larvae - ScienceDirect. Aquaculture. 234, 429–445.

Cejas, J.R., Almansa, E., Jérez, S., Bolaños, A., Felipe, B., Lorenzo, A., 2004. Changes in lipid class and fatty acid composition during development in white seabream (Diplodus sargus) eggs and larvae. Comparative Biochemistry and Physiology Part B: Biochemistry and Molecular Biology. 139, 209–216.

Churcher, A.M., Hubbard, P.C., Marques, J.P., Canario, A.V., Huertas, M., 2015. Deep sequencing of the olfactory epithelium reveals specific chemosensory receptors are expressed at sexual maturity in the European eel Anguilla anguilla. Mol Ecol. 24, 822–834.

Cowen, J. P., Holloway, C. F., 1996. Structural and chemical analysis of marine aggregates: In situ macrophotography and laser. Marine Biology.

Dekker, W., Casselman, J.M., 2014. The 2003 Québec declaration of concern about eel declines—11 years later: are eels climbing back up the slippery slope? Fisheries. 39, 613–614.

Folch, J., Lees, M., Sloane Stanley, G.H., 1957. A simple method for the isolation and purification of total lipides from animal tissues. J Biol Chem. 226, 497–509.

Gilani, G.S., Peace, R.W., 2005. Chromatographic determination of amino acids in foods. Journal of AOAC International. 88, 877–887.

Guedes, M., Costa-Pinto, A.R., Gonalves, V., Moreira-Silva, J., Neves, N.M., 2020. Sardine Roe as a Source of Lipids To Produce Liposomes. ACS Biomaterials Science and Engineering. XXXX.

Hsu, H.Y., Chen, S.H., Cha, Y.R., Tsukamoto, K., Lin, C.Y., Han, Y.S., 2015. De novo assembly of the whole transcriptome of the wild embryo, preleptocephalus, leptocephalus, and glass eel of Anguilla japonica and deciphering the digestive and absorptive capacities during early development. PloS one. 10, e0139105.

Huang, D., Liang, H., Ren, M., Ge, X., Ji, K., Yu, H., Maulu, S., 2021. Effects of dietary lysine levels on growth performance, whole body composition and gene expression related to glycometabolism and lipid metabolism in grass carp, Ctenopharyngodon idellus fry. Aquaculture. 530, 735806.

Ishikawa, S., Suzuki, K., Inagaki, T., Watanabe, S., Kimura, Y., Okamura, A., Otake, T., Mochioka, N., Suzuki, Y., Hasumoto, H., Oya, M., Miller, M., Lee, T.W., Fricke, H., Tsukamoto, K., 2001. Spawning time and place of the Japanese eel Anguilla japonica in the North Equatorial Current of the western North Pacific Ocean. Fisheries Science. 67, 1097–1103.

Jacoby, D., Gollock, M., 2014. The IUCN red list of threatened species, Version 2014. 3.

Jiang, T.B., Liu, L.P., Gao, X.Y., Chen, W.Y., Wu, J.M., 2012. The changes of serum biochemical components during carp pituitary extract and HCG induced maturation of the female Japanese eel (Anguilla japonica) (in Chinese). Journal of Fisheries of China. 36, 893–899.

Kleppe, L., Edvardsen, R.B., Furmanek, T., Taranger, G.L., Wargelius, A., 2014. Global transcriptome analysis identifies regulated transcripts and pathways activated during oogenesis and early embryogenesis in Atlantic cod. Molecular Reproduction & Development. 81, 619–635.

Koh, I.C.C., Hamada, D., Tsuji, Y., Okuda, D., Nomura, K., Tanaka, H., Ohta, H., 2017. Sperm cryopreservation of Japanese eel, Anguilla japonica. Aquaculture. 473, 487–492.

Kuan-Mei, H., Shingo, K., Yu-San, H., Aigo, T., Yoshiyuki, I., Caroline, D., 2018. Effect of ENSO events on larval and juvenile duration and transport of Japanese eel (Anguilla japonica). PloS one. 13, e0195544.

Kurokawa, T., Suzuki, T., Ohta, H., Kagawa, H., Unuma, T., 2002. Expression of pancreatic enzyme genes during the early larval stage of Japanese eel Anguilla japonica. Fisheries Science. 68, 736–744.

Langmead, B., 2012. Fast gapped-read alignment with bowtie 2. Nature Methods. 9, 357–359.

Li, B., Dewey, C.N., 2011. RSEM: accurate transcript quantification from rna-seq data with or without a reference genome. BMC Bioinformatics. 12, 1–16.

Li, C., Cheng, X.F., Hong, B., Chen, X.Y., Wu, Y.N., Li, H., 2016. Nutritional analysis and evaluation on eggs of Spinibarbus caldwelli. Chinese Journal of Animal Nutrition. 28, 2204–2212.

Lin, T.A., Xx, A., Hs, B., Jc, A., Chong, H.A., Hai, H.C., Hl, A., Yong, Z., Sl, A., 2020. Induction of oocyte maturation and changes in the biochemical composition, physiology and molecular biology of oocytes during maturation and hydration in the orange-spotted grouper (Epinephelus coioides). Aquaculture. 522.

Liu, L.P., Liu, D.P., Pu, J.C., Du, L., Chen, T.Y., Chen, W.Y., Zheng, C.J., Wu, X.F., Wu, J.M., 2017. Effects of, different initial diets on the survival and behavior characteristics of the larvae of Japanese eel (Anguilla japonica) (in Chinese). Journal of Fisheries of China. 41, 703–710.

Liu, Z.F., Gao, X.Q., Yu, J.X., Wang, Y.H., Guo, Z.L., 2018. Changes of protein and lipid contents, amino acid and fatty acid compositions in eggs and yolk-sac larvae of American shad (Alosa sapidissima). Journal of Ocean University of China. 17, 209–215.

Masuda, Y., Imaizumi, H., Usuki, H., Oda, K., Hashimoto, H., Teruya, K., 2012. Artificial completion of the Japanese eel, Anguilla japonica, life cycle: challenge to mass production. Bulletin of Fisheries Research Agency. 35, 111–117.

Matsushige, K., Yasutake, Y., Mochioka, N., 2019. Spatial distribution and habitat preferences of the Japanese eel, Anguilla japonica, at the reach and channel-unit scales in four rivers of Kagoshima Prefecture, Japan. Ichthyol Res. 67, 68–80.

Miller, M.J., Kimura, S., Friedland, K.D., Knights, B., Kim, H., Jellyman, D.J., Tsukamoto, K., 2009. Review of ocean-atmospheric factors in the Atlantic and Pacific oceans influencing spawning and recruitment of Anguillid eels. Marine, National Service, Fisheries. 69, 231–249.

Morrison, W.R., Smith, L.M., 1964. Preparation of fatty acid methyl esters and dimethylacetals from lipids with boron fluoride-methanol. J Lipid Res. 5, 600–608.

Mourente, G., Rosa, V., 1996. Changes in the content of total lipid, lipid classes and their fatty acids of developing eggs and unfed larvae of the Senegal sole, Solea senegalensis kaup. Fish Physiology & Biochemistry. 15, 221–235.

Ohkubo, N., Sawaguchi, S., Nomura, K., Tanaka, H., Matsubara, T., 2008. Utilization of free amino acids, yolk protein and lipids in developing eggs and yolk-sac larvae of Japanese eel Anguilla japonica. Aquaculture. 282, 130–137.

Okamura, A., Horie, N., Mikawa, N., Yamada, Y., Tsukamoto, K., 2014. Recent advances in artificial production of glass eels for conservation of anguillid eel populations. Ecol Freshw Fish. 23, 95–110.

Okamura, A., Yamada, Y., Horie, N., Mikawa, N., Tsukamoto, K., 2019. Long-term rearing of Japanese eel larvae using a liquid-type diet: food intake, survival and growth. Fisheries Science. 85, 687–694.

Okamura, A., Yamada, Y., Mikawa, N., Horie, N., Tsukamoto, K., 2020. Dietary supplementation with chitin hydrolysates for Anguilla japonica leptocephali. Fisheries Science. 86, 685–692.

Okamura, A., Yamada, Y., Horie, N., Mikawa, N., Tanaka, S., Kobayashi, H., Tsukamoto, K., 2013. Hen egg yolk and skinned krill as possible foods for rearing leptocephalus larvae of Anguilla japonica Temminck & Schlegel. Aquaculture Research. 44, 1531–1538.

Ozsolak, F., Milos, P.M., 2010. RNA sequencing: advances, challenges and opportunities. Nat Rev Genet. 12, 87–98.

Pfeiler, E., 1999. Developmental physiology of elopomorph leptocephali. Comparative Biochemistry & Physiology Part A Molecular & Integrative Physiology. 123, 113–128.

Politis, S.N., Sørensen, S.R., Mazurais, D., Servili, A., Zambonino-Infante, J.L., Miest, J.J., Clemmesen, C.M., Tomkiewicz, J., Butts, I.A., 2018. Molecular ontogeny of first-feeding European eel larvae. Frontiers in physiology. 9, 1477.

Pousão-Ferreira, P., Morais, S., Dores, E., Narciso, L., 1999. Eggs of gilthead seabream Sparus aurata L. as a potential enrichment product of Brachionus sp. in the larval rearing of gilthead seabream Sparus aurata L. Aquaculture Research. 30, 751–758.

R?Nnestad, I., Thorsen, A., Finn, R.N., 1999. Fish larval nutrition: a review of recent advances in the roles of amino acids. Aquaculture. 177, 201–216.

Robinson, M.D., Mccarthy, D.J., Smyth, G.K., 2010. EdgeR: a Bioconductor package for differential expression analysis of digital gene expression data. Bioinformatics. 26, 139–140.

Skoog, A., Alldredge, A., Passow, U., Dunne, J., Murray, J., 2008. Neutral aldoses as source indicators for marine snow. Marine Chemistry. 108, 195–206.

Tanaka, H., 2015. Progression in artificial seedling production of Japanese eel Anguilla japonica. Fisheries Science. 81, 11–19.

Tanaka, H., Kagawa, H., Ohta, H., 2001. Production of leptocephali of Japanese eel (Anguilla japonica) in captivity. Aquaculture. 201, 51–60.

Tanaka, H., Kagawa, H., Ohta, H., Unuma, T., Nomura, K., 2003. The first production of glass eel in captivity: fish reproductive physiology facilitates great progress in aquaculture. Fish Physiol Biochem. 28, 493–497.

Tsukamoto, K., 1992. Discovery of the spawning area for Japanese eel. Nature. 356, 789–791.

Wang, Z., Gerstein, M., Snyder, M., 2010. RNA-Seq : a revolutionary tool for transcriptomics. Nat Rev Genet. 10, 57–63.

Xiong, B.X., Long, L.Q., Su, X., Chang, Q., 1996. Crude protein and essential amino acid countens among eel feeds made in China (in Chinese). Journal of Huazhong Agricultural University. 15, 60–63.

Yanes-Roca, C., Rhody, N., Nystrom, M., Main, K.L., 2009. Effects of fatty acid composition and spawning season patterns on egg quality and larval survival in common snook (Centropomus undecimalis). Aquaculture. 287, 335–340.

Yang, J.J., Jiang, Z.Q., Zuo, R.T., Wang, S.Y., Wen, S.H., Sun, H., 2014. Nutritional analysis and evaluation on eggs of Hemitripterus villosus (in Chinese). Chinese Journal of Animal Nutrition. 26, 1103–1110.

Youqing, X., Weifeng, L., Zhaokun, D., 2010. Ef fects of Polyunsaturated Fatty Acids on Immunity and Survival of Fish and Their Mechanisms. Chinese Journal of Ani mal Nutrition. 22, 551-556(in Chinese).

Zheng, T.T., Zhou, J., Weng, X., Chen, L.J., Cheng, W.J., Pang, J., Liang, P., 2020. Analysis and evaluation of nutritional components of four types of marine aquatic roes. Food and Fermentation Industries. 46, 244–249.

Zhou, L.L., Zhu, S.X., Li, X.X., Dan, X.M., Liu, L., Pan, Q., Li, G.F., Gan, L., 2014. Basic components analysis and nutritive value evaluation of Anguilla bicolor pacifica muscle. Guangdong Agricultural Sciences. 41, 105–109.

Zhou, X.Q., Zhao, C.R., Jiang, J., Feng, L., Liu, Y., 2008. Dietary lysine requirement of juvenile Jian carp (Cyprinus carpiovar. Jian). Aquaculture Nutrition. 14, 381–386.

